# Blood Omics Models for System-Specific Mortality Risk Estimation

**DOI:** 10.1101/2024.11.27.625723

**Authors:** Matías Fuentealba, Thalida Em Arpawong, Kevin Schneider, Eileen M Crimmins, David Furman

## Abstract

Traditional blood-based aging clocks provide an estimate of a person’s overall biological age. However, physiological systems and organs age at different rates in an individual, and anti-aging interventions often target specific physiological systems. Therefore, there is a growing need for methods capable of assessing biological age at the level of specific physiological systems. Here, we used blood chemistry and cell count data from 456,180 individuals in the UK Biobank (UKB) to develop mortality-based predictors of biological age across 9 physiological systems matching WHO’s International Classification of Disease (ICD-10) chapters (DiseaseAge). We applied DiseaseAge to the Health and Retirement Study (HRS) cohort and validated its ability to identify biologically older systems in individuals diagnosed or deceased from age-related diseases affecting those systems. For instance, individuals diagnosed with high blood pressure, heart attack, congestive heart failure, or angina exhibited a biologically older circulatory system than other systems. Similarly, individuals with accelerated aging in the circulatory, musculoskeletal, or respiratory systems displayed higher risk of mortality from conditions associated with these systems. Additionally, we showed that individuals within the top 5% biologically older metabolic, circulatory, respiratory and mental systems exhibited increased risk of developing diabetes, high blood pressure, lung disease and dementia, respectively. Finally, we used metabolomics and proteomics data in the UKB and epigenomics and transcriptomics in HRS to generate omics surrogates of DiseaseAge for all physiological systems and created an online resource for their calculation.

## Main

Quantifying biological age is a cornerstone of aging research as it provides a framework for assessing aging rates in individuals and it enables the identification of interventions to improve health during aging. While survival is the gold standard for assessing aging effects in model organisms^1^, it is impractical in humans as it requires clinical trials spanning decades. Since 2013, following the publication of the Hannum^2^ and Horvath clocks^3^ epigenetic aging clocks have become the standard for assessing biological age in living humans, enabling the evaluation of anti-aging interventions within a reasonable timeframe. Since then, other epigenetic clocks have been developed to estimate biological age^4–6^ and track the rate of aging^7^, in some cases using sophisticated machine learning methods^8,9^ or different omics such as transcriptomics^10,11^, proteomics^12^, and metabolomics^13^. Despite this great deal of progress, a common limitation of omics clocks is that they provide an estimate of the biological age of the whole body and different physiological systems are known to age at different rates with biologically older systems being more prone to diseases. Also, health interventions and disease treatments often target specific physiological systems. Thus, new omics clocks designed to estimate biological age in specific physiological systems have emerged as a means to identity faster-aging systems and guide targeted interventions for improvement. For example, Sehgal et al. (2023)^14^ selected clinical biomarkers associated with 11 different physiological systems to build a predictive model of all-cause mortality using epigenetics. They showed that system-specific scores were significantly associated with meaningful disease outcomes, such as the blood system score predicting time-to-leukemia and the heart system score predicting time-to-coronary heart disease. In another study, Wyss-Coray’s group^15^ identified genes whose expression is elevated in each organ and then used protein abundance levels of these genes in the blood to generate a predictors of chronological age for each organ. They showed that 18% of people exhibited accelerated aging in at least one organ, with only 1.7% of participants showing accelerated aging in multiple organs. More recently, Reicher et al. (2024)^16^ categorized clinical, physiological and behavioral parameters to predict the biological age associated with 14 physiological systems. Interestingly, they found sex-specific patterns in biological aging which were strongly associated with a higher prevalence of age-related conditions. Although these methods represent a significant improvement over traditional blood-based clocks in assessing system-specific aging, they still have limitations such as reliance on manual feature selection prone to research bias, limited cross-cohort validation, and a focus on single omics, which reduces applicability.

Here, we used blood biochemistry and cell count data from 456,180 individuals in the UK Biobank and elastic net Cox penalized regression to predict cause-specific mortality risk associated with diseases grouped into 9 physiological systems based on the International Classification of Diseases (ICD-10): circulatory, digestive, genitourinary, infectious, mental, metabolic, musculoskeletal, nervous and respiratory (Sup. Table 1). Additionally, we created a predictive models of all-cause mortality risk. To improve interpretability, we transformed the mortality risk estimations into age values by matching the distribution of mortality risk and chronological age, resulting in an estimation for each physiological system termed DiseaseAge. DiseaseAge displays moderate to high correlations with chronological age across all physiological systems, with the smallest correlations observed in the metabolic (R = 0.54) and digestive (R = 0.61) systems, and the highest in the mental (R = 0.96) and nervous (R = 0.93) systems (Fig. 1a). We assessed the performance of predicting mortality risk using the concordance index (C-index) and observed that all system-specific mortality predictors outperformed all-cause mortality models (C = 0.78 to 0.89) (Fig. 1b). Interestingly, there was no correspondence between the correlation of DiseaseAge with chronological age, and the C-index for system-specific mortality, which indicates that more accurate estimators of chronological aging are not necessarily better predictors of health (R = -0.19, p = 0.58).

**Fig. 1.**
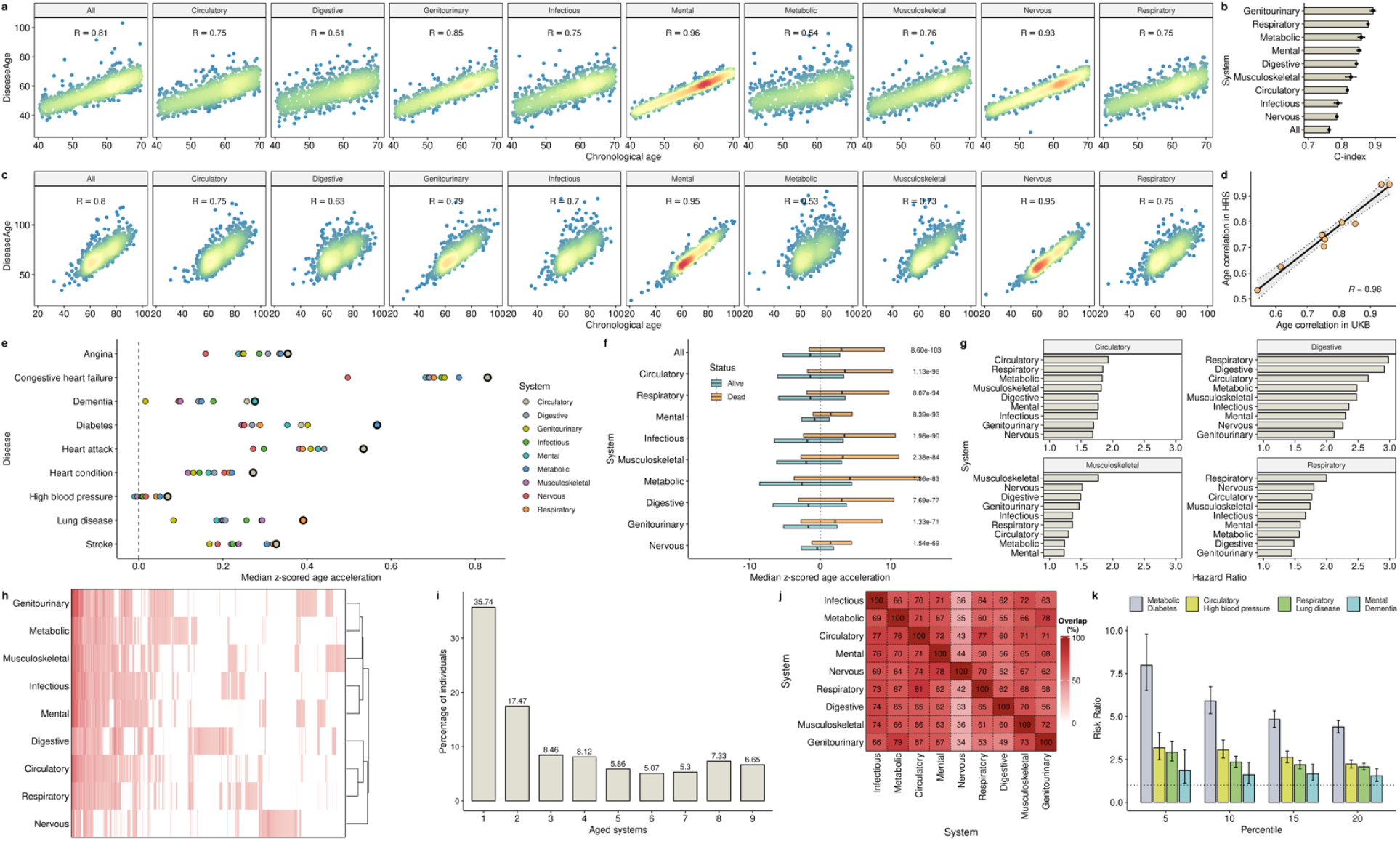
DiseaseAge Predicts Biological Aging and Mortality Across Physiological Systems. a, Correlation between chronological age and DiseaseAge across physiological systems. Dot colors indicate the sample density. b, Predictive performance of system-specific and all-cause mortality models of DiseaseAge. c, Validation of DiseaseAge in the HRS cohort. d, Dependency of the correlations between DiseaseAge and chronological age in UKB and HRS. e, Accelerated aging for each system in individuals diagnosed with age-related diseases. Highlighted circles indicate systems associated with the disease analyzed. f, All-cause DiseaseAge differences between living and deceased individuals. g, Cause-specific mortality risk associated with accelerated aging in different systems. h, Individuals with extreme accelerated aging in at least one physiological system. i, Percentage of individuals with extreme accelerated aging across multiple systems. j, Co-occurrence of accelerated aging across physiological systems. It indicates the percentage of individuals with accelerated aging in the systems in the columns that also displayed accelerated aging in the systems in the rows. k, Association between extreme accelerated aging and age-related disease risk.

To enable cross-cohort validation, the DiseaseAge models were built using a set of features shared between UKB and HRS cohorts (Sup. Table 2), and all down-stream analysis were performed in HRS. In HRS, DiseaseAge results in strikingly similar correlation with chronological age across systems (correlation of age correlations = 0.98, Fig. 1c-d). As initial validation in HRS, we evaluated whether individuals diagnosed with age-related disease exhibited greater accelerated aging in the corresponding physiological system. Notably, we observed that for all the diseases analyzed, the corresponding physiological system displayed accelerated aging (Fig. 1e). For example, individuals diagnosed with angina, congestive heart failure, heart attack, heart condition, high blood pressure, and stroke displayed higher accelerated aging in the circulatory system. Similarly, individuals with dementia, diabetes and lung disease showed accelerated aging in the mental, metabolic, and respiratory systems, respectively. As further validation of the specificity of the model, we tested if accelerated aging was associated with an increased risk of death from conditions associated with different systems. We first compared living and deceased individuals (due to any cause) and observed that all-cause DiseaseAge resulted in the strongest biological age differences (p = 8.6e-103, Fig. 1f). We then examined cause-specific mortality and observed that individuals with accelerated aging in the circulatory, musculoskeletal or respiratory systems, had a higher risk of death from conditions in those systems (Fig. g) more than other systems.

Next, we sought to identify individuals with extreme phenotypes of accelerated aging in specific systems. We identified 887 individuals (9.4%) with accelerated aging, defined as exceeding two standard deviations above the mean, in at least one system (Fig. 1h). Interestingly, 35% these individuals (n = 317) displayed extreme accelerated aging in a single system, while only 6.6% (n = 59) had extreme accelerated aging across all physiological systems (Fig. 1i). Among the 570 individuals with extreme accelerated aging in more than one system, we examined which systems tended to co-occur more often. We found that 81% of the individuals with accelerated aging in the circulatory systems also showed accelerated aging in the respiratory system (Fig. 1j). Interestingly, accelerated aging in the nervous system typically did not co-occur with accelerated aging in other systems. We also analyzed if extreme accelerated aging was associated with a higher risk of age-related diseases. We found that individuals among the 5% highest accelerated aging in the metabolic system displayed a 7.9-fold increase in the risk of diabetes (p = 4.04e-109) (Fig. 1k). Similarly, extreme accelerated aging in the circulatory, respiratory, and mental systems was associated with significantly higher risk of high blood pressure (RR = 3.17, p = 2.27e-25), lung disease (RR = 2.92, p = 9.69e-23) and dementia (RR = 1.85, p = 2.43e-02), respectively.

Finally, we explored whether molecular changes detectable in omics data could improve the sensitivity of the predictors to system-specific mortality risk, as it has been previously observed^4,5^. To test this, we used elastic net regression to build omics-based surrogates of DiseaseAge using proteomics and metabolomics data in the UKB and epigenomics and transcriptomics data in HRS (Fig. 2a). Overall, we observed a strong correlation between DiseaseAge and its omics-based across physiological systems. On average, the highest correlations were observed when using proteomics (average R = 0.91), followed by epigenomics (average R = 0.81) data to predict DiseaseAge. Interestingly, metabolomics data was particularly effective at predicting Metabolic DiseaseAge in comparison with other systems (R = 0.73). Also, Mental and Nervous DiseaseAge were better predicted with Epigenomics than other systems (R = 0.89 and 0.9), while Digestive DiseaseAge was not well predicted with either Epigenomics or Transcriptomics. We then compared the performance of predicting cause-specific mortality of DiseaseAge and the omics surrogates in the UKB and HRS (Fig. 2b). Notably, we found that proteomics-based DiseaseAge consistently outperformed the original DiseaseAge in predicting mortality across all physiological systems, while metabolomic, epigenetic and transcriptomic-based estimates performed worse but similar to DiseaseAge in predicting cause-specific mortality. To facilitate the calculation of DiseaseAge from any type of omics data, we next created an online resource (https://diseaseage.furmanlab.org). The application requires as input a matrix in tab- or comma-separated values, with the first column containing feature names and the remaining columns containing sample values. The results include the estimation of DiseaseAge for each physiological system in every sample and the coverage of the model features in the input data.

**Fig. 2.**
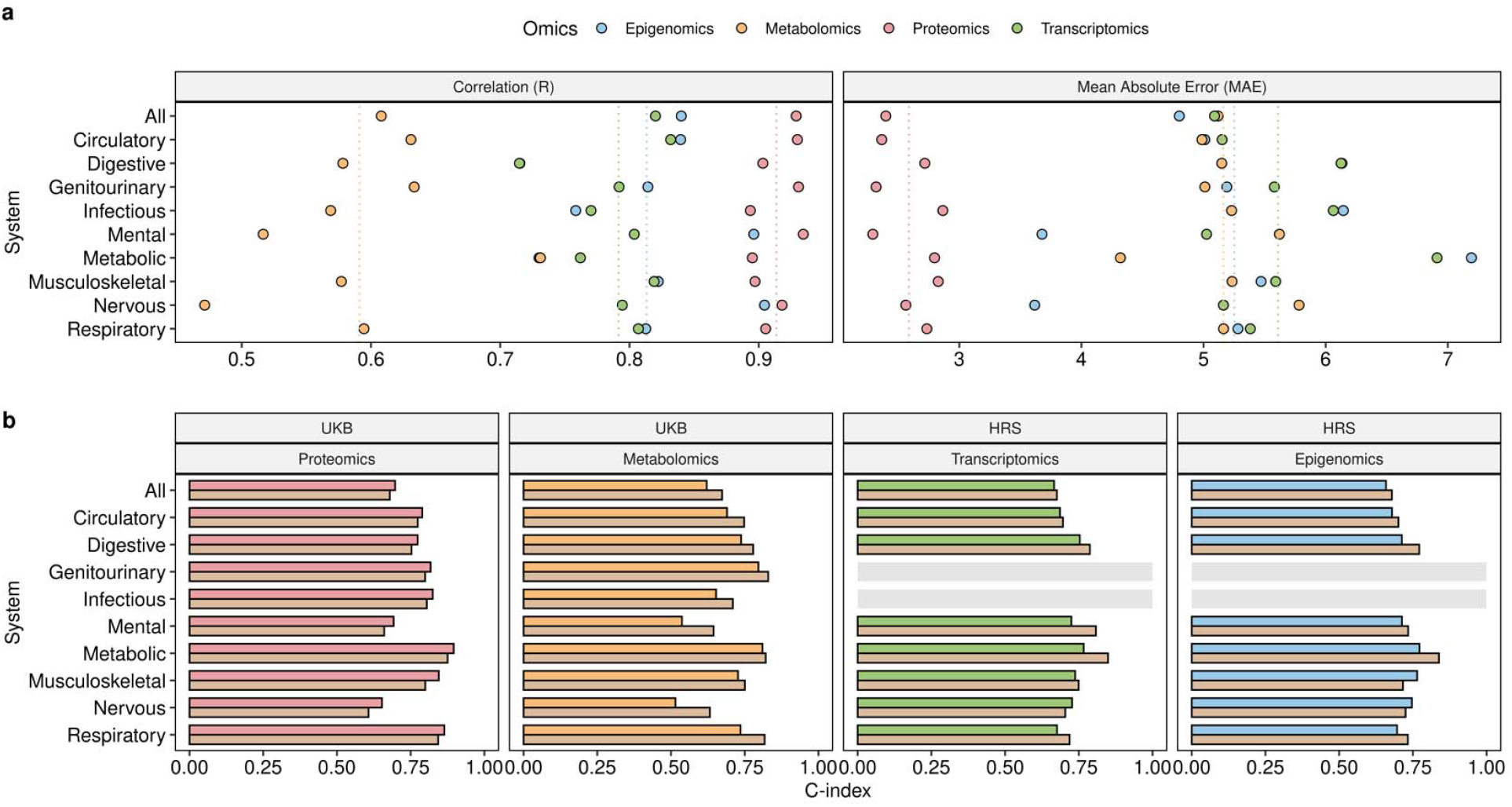
Omics-based surrogates of DiseaseAge and their performance in predicting system-specific mortality. a, Correlation and error between DiseaseAge and omics-based surrogates built using proteomics and metabolomics data in the UKB and epigenomics and transcriptomics data in the HRS. b, Performance of DiseaseAge and omics-based surrogates in predicting system-specific mortality. No deaths from genitourinary or infections were reported and coded in HRS. Light brown bars indicate the estimates from DiseaseAge.

## Discussion

Here, we developed a system-specific predictor of biological age based on mortality data. Considering that only 8.8% of UKB participants have experienced mortality, we anticipate that future iterations of the models will improve in performance and coverage of systems as more data become available in the coming years. Additionally, given sample limitations, we have only used the top level of the ICD-10 hierarchy to describe physiological systems. However, the ICD-10 coding system has the potential to generate predictors at higher levels of resolution enabling the evaluation of different organ functions or structures associated with specific groups of diseases. In comparison with other system-specific predictors of biological age^14,16,17^, our approach is the first to use a data-driven approach to identify features predictive of each system’s function, making the models less prone to exclude features relevant for the physiological system compared to manual selection. Through cross-cohort calculations we validated the model’s specificity by observing greater accelerated aging in systems consistent with the diseases diagnosed or causes of death, and an increased risk for specific diseases in individuals with extreme accelerated aging in the corresponding systems. Finally, we also created multiple omics surrogates of our system-specific aging predictors, enabling researchers to evaluate the effects of diseases and interventions in each physiological system using either epigenomics, transcriptomics, proteomics or metabolomics. Compared to previous methods designed using epigenomics or proteomics data^14,15^, these models provide broader applicability and cross-omics comparisons.

## Methods

### Blood biochemistry and blood count data

Data were extracted from UKB categories 100081 and 17518 respectively. We considered only measurements that were also present in HRS (Sup. Table 2). Among the 38 features obtained measured in 502,175, we excluded those with less than 80% of the features measured resulting in 456,180 samples. Missing values were imputed using the nearest neighbor average method implemented in the R package impute^18^. In HRS, blood biochemistry and blood count data were obtained from the 2016 Venous Blood Study (VBS) Version 2 and VBS 2016 Additional Assays data. Age and sex were obtained from the Cross-Wave Tracker File Early 2022 V1.0. Measurements were converted to match the units used in the UKB. From the 9933 individuals in HRS, 357 were removed due to missing values in more than 20% of the 38 features. The remaining values were imputed using nearest neighbor average.

### Survival data

In the UKB, time to death was calculated as the difference between age at death (field 40007) and age when attending assessment center (field 21003). In living individuals, it corresponded to the difference between the date of attending the assessment center (field 53) and date when last seen. Causes of death as International Classification of Diseases (ICD-10) codes were obtained using data field 40001. In HRS, time to death was calculated as the difference between the date when blood was given and the date of death (fields KNOWNDECEASEDYR, KNOWNDECEASEDMO). In living individuals, it corresponded to the difference between last seen alive (fields LASTALIVEYR, LASTALIVEMO) and the date when blood was given. Causes of death were obtained from the 2020 HRS Exit questionnaire (field XRA133M1M). Although the cause of death in HRS were not coded directly as ICD-10, the coding system was equivalent.

### DiseaseAge calculation

To improve the normality of the features, we calculated kurtosis and skewness of the features before and after log transformation and observed that 25 of the features (excluding age and sex), showed reduction after the transformation (Sup. Table 3). These transformed features along with the remaining untransformed features were used for modeling. Causes of death were grouped into system-level categories, representing different physiological systems. We only kept 9 systems with at least 100 deaths associated. For each system, we performed 100 iterations of elastic net cox penalized regression with glmnet^19^ using the blood biochemistry and cell count data as input to predict the mortality risk. In each iteration, diseased individuals were compared against a random set of controls composed of 10 individuals for each dead individual. This procedure was designed to reduce the unbalanced ratio of dead and alive individuals while maximizing statistical power. In each model, the lambda parameter was selected to maximize C-index. To enhance model robustness and specificity, we identified the features that were selected in all 100 iterations and performed an unpenalized cox regression model using 10-fold cross validation. Mortality risk estimations were transformed into age values by matching the distribution of mortality risk and chronological age as previously described^5^. Using cross-validated predictions we calculated the C-index of each model using the function survConcordance in the survival R package^20^.

### Disease association and cause-specific mortality

Using the 2016 HRS Core data we identified individuals diagnosed with high blood pressure (PC005), diabetes (PC010), lung disease (PC030), heart condition (PC036), heart attack (PC040), angina (PC045), congestive heart failure (PC048), stroke (PC053) and dementia (PC273). In each system we calculated the accelerated aging of each individual as the residual between a linear regression between DiseaseAge and chronological age. We then scaled (z-transform) the values to make them comparable across different systems. For each condition, we calculated the median scaled accelerated aging of individuals using the models for the 9 different systems. Using the scaled accelerated aging of each individual in each system, we compared the values between dead and alive individuals using a t-test. In the case of cause-specific mortality we used the scaled accelerated aging to fit a proportional hazard regression model of dead due specific causes including cancer, and circulatory, respiratory and digestive conditions. Then we compared the hazard ratios associated with the scaled accelerated aging in each system.

### Extreme accelerated aging

We considered individuals with extreme accelerated aging if they had a scaled accelerated aging above two (standard deviation units). Overlaps between individuals with extreme accelerated aging in different systems were calculated based on the percentage of individuals with extreme accelerated aging in one system that also showed extreme accelerated aging in another system. To calculate the disease risk associated with extreme accelerated aging we calculated the risk of diseases in individuals within the 5%, 10%, 15% and 20% highest accelerated aging and individuals with scaled accelerated aging below two in all systems (i.e. remaining samples). The risk ratio was calculated using the function riskratio implemented in the R package epitools^21^.

### Omics data processing

Epigenomics (HRS): DNA methylation was measured using DNA extracted from the buffy coat using the Infinium Methylation EPIC BeadChip by the Advanced Research and Diagnostics Laboratory at the University of Minnesota. DNA samples were randomized across analytic plates by age, cohort, sex, education, and race/ethnicity along with 39 pairs of blinded duplicates. Analysis of duplicate samples showed a correlation > 0.97 across all CpG sites. Data preprocessing and quality control were performed using the minfi package in R^22^. A total of 3.4% of the methylation probes were removed from the final data because their detection p-value fell below the threshold of 0.01 (n = 29,431 out of 866,091). Analysis for failed samples was done after removal of detection p-value failed probes. A total of 58 samples were removed after applying a 5% cutoff. Sex-mismatched samples and any controls (cell lines, blinded duplicates) were also not included in the analyses. The final analytic data set includes 97.9% (n = 3,921) of the originally plated samples. Missing beta methylation values were imputed with the median beta methylation value of the given probe across all samples^23^.

Transcriptomics (HRS): RNA was extracted from whole blood stored in Paxgene tubes by using the Paxgene Blood miRNA Kit. Extracted RNA is then stored at -80°C until further analysis. Total RNA isolates were quantified using a fluorimetric RiboGreen assay. Total RNA samples were treated with the Globin-Zero Gold rRNA Removal Kit (Illumina Inc.) to deplete ribosomal RNA and globin prior to creating sequencing libraries using Illumina’s stranded mRNA Sample Preparation kit (Cat. # RS-122-2101). One microgram of total RNA was oligo-dT purified using oligo-dT coated magnetic beads, fragmented and then reverse transcribed into cDNA, fragmented, blunt-ended, and ligated to indexed (barcoded) adaptors and amplified using 15 cycles of PCR. Indexed libraries were then normalized, pooled and size selected to 320bp +/-5% using Caliper’s XT instrument. Samples were sequenced using 2*50 bp paired-end reads to a minimum of 20 million reads per sample on NovaSeq at the University of Minnesota Genomics Center. All samples were processed through the HRS RNAseq QC analysis pipeline at the University of Minnesota, which is an extended version of the TopMed/GTEX analysis pipeline (https://github.com/broadinstitute/gtex-pipeline/blob/master/TOPMed_RNAseq_pipeline.md). The STAR aligner was used for alignment of the sequence reads to the GRCh38 human reference genome along with GENCODE 30 annotations^24^. All quality control analyses were performed using an updated version of RNASeQC 2.3.4^25^ and estimated quality control metrics to obtain the final data (n = 3,658). The read counts from each sample were combined into a count file. The package edgeR^26^ calcNormFactors function and RLE (relative log expression) normalization was used to account for compositional differences between the libraries. RLE is the scaling factor method in which the median library is calculated from the geometric mean of all columns and the median ratio of each sample to the median library is taken as the scale factor. Then the cpm() function in edgeR was implemented on the normalized DGEList object to estimate the log2 counts-per-million (log2cpm) with a prior.count=1.

Metabolomics (UKB): NMR metabolomics was performed by Nightingale Health Plc. on EDTA plasma samples (Nightingale Health Metabolic Biomarkers Companion document. https://biobank.ndph.ox.ac.uk/showcase/refer.cgi?id=130. Accessed 22 Nov 2024). Data from phase 1 and phase 2 of the study were included in the data request from the UK Biobank Showcase and include absolute concentration measurements for 168 metabolites, 81 metabolite ratios and 2 derived variables. Measurements were completed between June 2019 and April 2020 (phase 1) and April 2020 and June 2022 (phase 2). All processed values provided by the UK Biobank have either passed quality control or have been determined that the quantification of the biomarker is valid.

Proteomics (UKB): The Olink target platform panel dataset was downloaded from the UK BioBank Table Exporter application on the Research Analysis Platform for all proteins in the panel from EDTA plasma samples for 53,014 participants at their baseline visit between 2006 and 2010. Protein values are provided by the UK Biobank as normalized protein expression (NPX) values, which are calculated within and between each batch to the reference batch (batch 1). Within batch normalization is calculated for all plates within a batch by subtracting the log2 ratio of counts of the sample to counts of the extension control by the protein-specific median value of the plate controls and for each sample subtracting the protein-specific median value from the plate (UK Biobank Team. UKB – Olink Explore 1536 -Data Normalization Strategy. https://biobank.ndph.ox.ac.uk/showcase/refer.cgi?id=4656. Accessed 22 Nov 2024.). Median NPX values from batch 1 were used to calculate adjustment factors for between batch normalization.

### Omics clocks

Omics clocks were trained using omics data and a 10-fold cross-validated elastic net regression using glmnet^19^ to predict DiseaseAge. Lambda values were selected to minimize mean absolute error. We calculated the concordance index using DiseageAge or the predicted DiseaseAge using omics from the cross-validation using the survival R package^20^. To maximize the number of deceased individuals we used records during 2018 and 2020 HRS exit data (fields XQA133M1M, XQA133M2M, XRA133M1M, XRA133M2M). The web server to perform the calculations of omics DiseaseAge was generated using the R package shiny^27^.

## Notes

### Competing Interest Statement

The authors have declared no competing interest.

